# N-glycolylneuraminic acid binding of avian H7 influenza A viruses

**DOI:** 10.1101/2020.12.21.423767

**Authors:** Cindy M. Spruit, Xueyong Zhu, Frederik Broszeit, Alvin X. Han, Roosmarijn van der Woude, Kim M. Bouwman, Michel M. T. Luu, Colin A. Russell, Ian A. Wilson, Geert-Jan Boons, Robert P. de Vries

## Abstract

Influenza A viruses initiate infection by binding to glycans with terminal sialic acids present on the cell surface. Hosts of influenza A viruses variably express two major forms of sialic acid, N-acetylneuraminic acid (NeuAc) and N-glycolylneuraminic acid (NeuGc). NeuGc is produced in the majority of mammals including horses, pigs, and mice, but is absent in humans, ferrets, and birds. Intriguingly, the only known naturally occurring influenza A viruses that exclusively bind NeuGc are the extinct highly pathogenic equine H7N7 viruses. We determined the crystal structure of a representative equine H7 hemagglutinin (HA) in complex with its NeuGc ligand and observed a high similarity in the receptor-binding domain with an avian H7 HA. To determine the molecular basis for NeuAc and NeuGc specificity, we performed systematic mutational analyses, based on the structural insights, on two distant avian H7 HAs. We found that mutation A135E is key for binding α2,3-linked NeuGc but does not abolish NeuAc binding. Interestingly, additional mutations S128T, I130V, or a combination of T189A and K193R, converted from NeuAc to NeuGc specificity as determined by glycan microarrays. However, specific binding to NeuGc-terminal glycans on our glycan array did not always correspond with full NeuGc specificity on chicken and equine erythrocytes and tracheal epithelium sections. Phylogenetic analysis of avian and equine H7 HAs that investigated the amino acids at positions 128, 130, 135, 189, and 193 reveals a clear distinction between equine and avian residues. The highest variability in amino acids (four different residues) is observed at key position 135, of which only the equine glutamic acid leads to binding of NeuGc. The results demonstrate that avian H7 viruses, although genetically distinct from equine H7 viruses, can bind NeuGc after the introduction of two to three mutations, providing insights into the adaptation of H7 viruses to NeuGc receptors.

**Author summary:** Influenza A viruses cause millions of cases of severe illness and deaths annually. To initiate infection and replicate, the virus first needs to bind to a structure on the cell surface, like a key fitting in a lock. For influenza A virus, these ‘keys’ (receptors) on the cell surface are chains of sugar molecules (glycans). The terminal sugar on these glycans is often either N-acetylneuraminic acid (NeuAc) or N-glycolylneuraminic acid (NeuGc). Most influenza A viruses bind NeuAc, but a small minority binds NeuGc. NeuGc is present in species like horses, pigs, and mice, but not in humans, ferrets, and birds. Therefore, NeuGc binding could be a determinant of an Influenza A virus species barrier. Here, we investigated the molecular determinants of NeuGc specificity and the origin of viruses that bind NeuGc.

## Introduction

Influenza A viruses can infect a broad range of animals, including mammalian and avian species. Infection is initiated when the hemagglutinin (HA) on the outside of a virus particle binds to glycans with terminal sialic acid on the cell surface. The vast majority of Influenza A viruses use a glycan with a terminal N-acetylneuraminic acid (NeuAc) as their receptor, although some strains use N-glycolylneuraminic acid (NeuGc) instead. Sialic acids are bound in the receptor-binding site (RBS) of the HA, consisting of conserved residues (Y98, W153, H183, and Y195) and structural features (130-, 150-, and 220-loop and 190-helix) [1]. Amino acid mutations in or near the RBS can change HA binding specificity, as shown extensively for HAs binding to either α2,3-linked or α2,6-linked NeuAc [2-6].

The ability of viruses to bind either α2,3-linked or α2,6-linked sialic acids is a host determinant. Binding to either NeuAc or NeuGc could likewise affect the host range. NeuGc is only present in species that express an active form of the enzyme cytidine monophosphate CMP-N-acetyl neuraminic acid hydroxylase (CMAH), which facilitates the hydroxylation of NeuAc to convert it to NeuGc. The gene encoding CMAH, mainly expressed in mammalian species, has been partially or completely lost at several distinct events during evolution [7], causing NeuGc to be absent in, among others, humans, ferrets, European dogs, and avian species [8-10].

In species that generate NeuGc, its percentage of the total sialic acid content varies. For instance, pig trachea contains an equal amount of NeuAc and NeuGc, while 90% of the sialic acids on equine trachea and erythrocytes is NeuGc [11-14]. The high NeuGc content in horses may explain why equine H7N7 viruses are the only known influenza A viruses that bind α2,3-linked NeuGc [15,16]. Highly pathogenic equine H7 viruses have not been isolated since 1978 and are, therefore, thought to be extinct [17,18]. Currently, horses are mainly infected by circulating H3N8 influenza viruses and bind NeuAc instead. Unlike equine H7 strains, avian and human H7 viruses bind NeuAc [16]. It is still unclear where equine H7 viruses originated from and what the molecular determinants of NeuGc specificity are.

Here, we investigated receptor binding specificities of avian H7 HAs to identify the origin of equine H7 viruses. Inspired by the crystal structure of the equine H7 HA in complex with its ligand NeuGc, we performed targeted mutagenesis of avian H7 viruses. Several combinations of mutations were found that enabled avian H7 viruses to bind NeuGc. Our results demonstrate a phenotypical relationship between avian and equine H7 viruses despite their substantial genetic distance.

## Results

### Crystal structure of an equine H7 HA in complex with receptor analog 3′-GcLN and its similarity to an avian H7 HA

We previously reported the crystal structure of the HA of A/Equine/New York/49/73 H7N7 (H7eq) without a ligand (PDB: 6N5A [15]). To understand the structural basis for NeuGc specificity of H7eq, we determined the crystal structure of H7eq in complex with its natural ligand 3′-GcLN (NeuGcα2-3Galβ1-4GlcNAc) at 2.05 Å resolution (Fig 1A and S1 Table). The electron density for the 3′-GcLN ligand could be fitted well for all three monosaccharides (Fig 1B). H7eq binds 3′-GcLN mainly through NeuGc-1, but the interactions extend over the 220-loop with hydrogen bonds between Gal-2 and the main-chain carbonyl oxygen of G225 and between GlcNAc-3 and the side chain of Q222 (Fig 1A).

**Fig 1.**
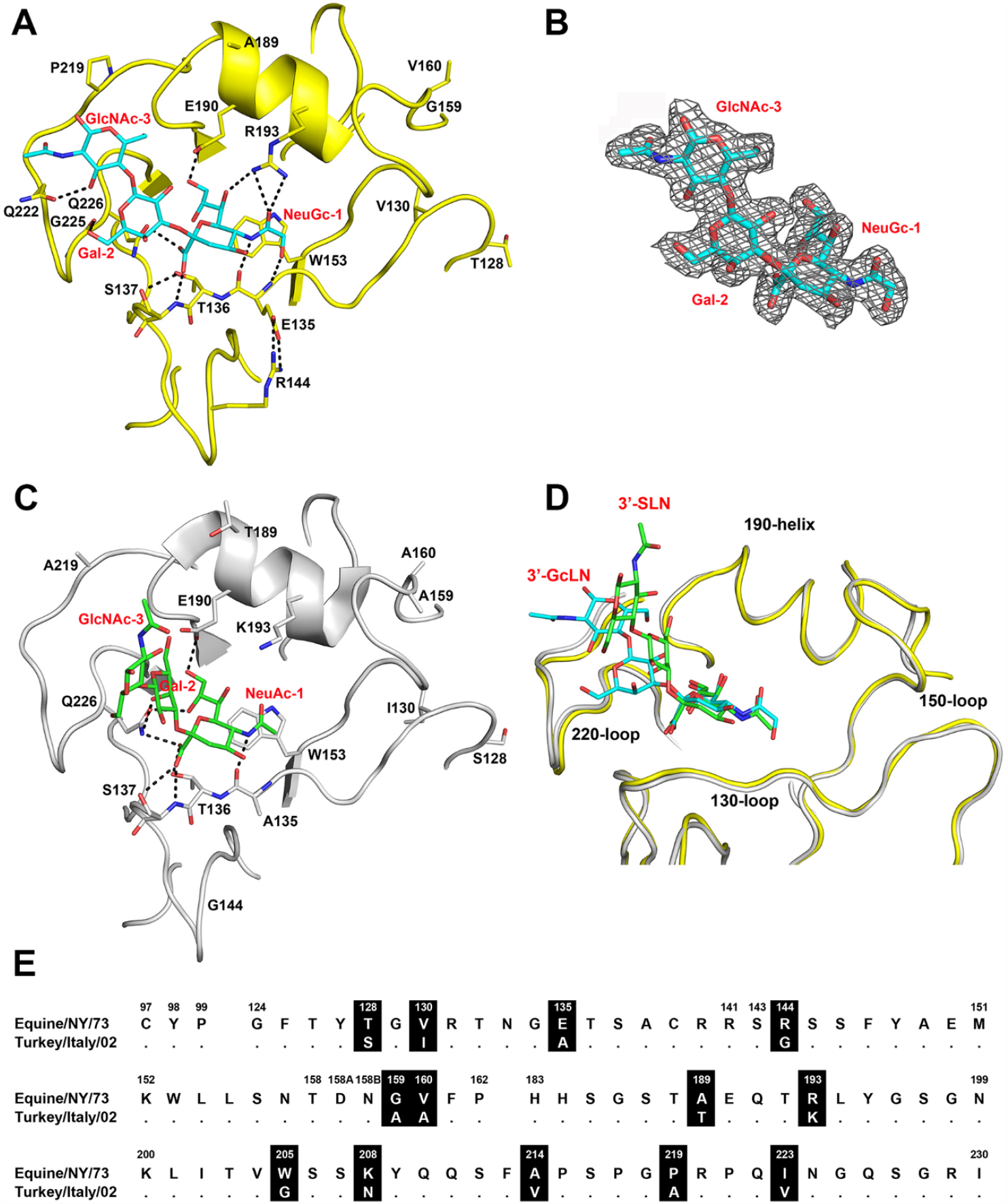
Comparison of the HA of A/Equine/New York/43/73 H7N7 and A/Turkey/Italy/214845/03 H7N3. (**A**) RBS structure of H7eq (yellow) in complex with 3′-GcLN (NeuGcα2-3Galβ1-4GlcNAc, cyan). (**B**) Electron density 2Fo-Fc map at 1σ level for receptor analog 3′-GcLN. (**C**) RBS structure of H7tu (grey) in complex with 3′-SLN (NeuAcα2-3Galβ1-4GlcNAc, green) (PDB: 4BSI). (**D**) Superimposition of the RBS structures of H7eq and H7tu and their ligands 3′-GcLN and 3′-SLN. The coloring scheme is following A and C. (**E**) Alignment of the RBS residues of H7eq and H7tu, with amino acid positions (H3 numbering) indicated above the alignment, non-conserved residues highlighted in black, and dots indicating identical amino acids.

The RBS structures of H7eq and A/Turkey/Italy/214845/02 H7N3 (H7tu) (PDB: 4SBI, [20] appear to be very similar (Fig 1C-D), although the turkey strain binds NeuAc instead of NeuGc and was isolated almost 30 years after the equine strain. Nevertheless, 85% of HA1 residues are identical and the amino acid sequences around the RBS differ at 13 positions (Fig 1E and S1). The NeuGc-Gal bond of 3′-GcLN in the H7eq complex adopts a *cis* conformation, which is consistent with our previous findings in the structure of 3′-GcLN in complex with the A/Vietnam/1203/2004 H5N1 Y161A mutant that shifts receptor specificity from NeuAc to NeuGc [15]. On the contrary, the NeuAc-Gal bond in the avian receptor analog 3′-SLN (NeuAcα2-3Galβ1-4GlcNAc) in complex with H7tu adopts a *trans* NeuAc-Gal bond (Fig 1C-D).

In the H7eq 3′-GcLN structure, the 1-hydroxyl group of NeuGc-1 forms a hydrogen bond with the main-chain nitrogen of E135 and the E135 side chain forms a salt bridge with R144 (Fig 1A). The amino acid at position 193 is known to be an important determinant of receptor specificity [21-24]. In the crystal structure of H7eq and 3′-GcLN, R193 forms a hydrogen bond with NeuGc-1 (Fig 1A). In comparison, K193 in H7tu, with a shorter side-chain, is not in hydrogen bond distance with the NeuAc-1 of 3′-SLN (Fig 1C).

Despite the similarities in RBS structures, we found that H7tu binds solely to α2,3-linked NeuAc on the glycan array (Fig 2B), whereas H7eq exclusively binds to α2,3-linked NeuGc [15]. To decipher which residues determine NeuGc and NeuAc receptor specificity, targeted mutagenesis was performed on H7tu by replacing residues in the RBS with H7eq-like amino acids.

**Fig 2.**
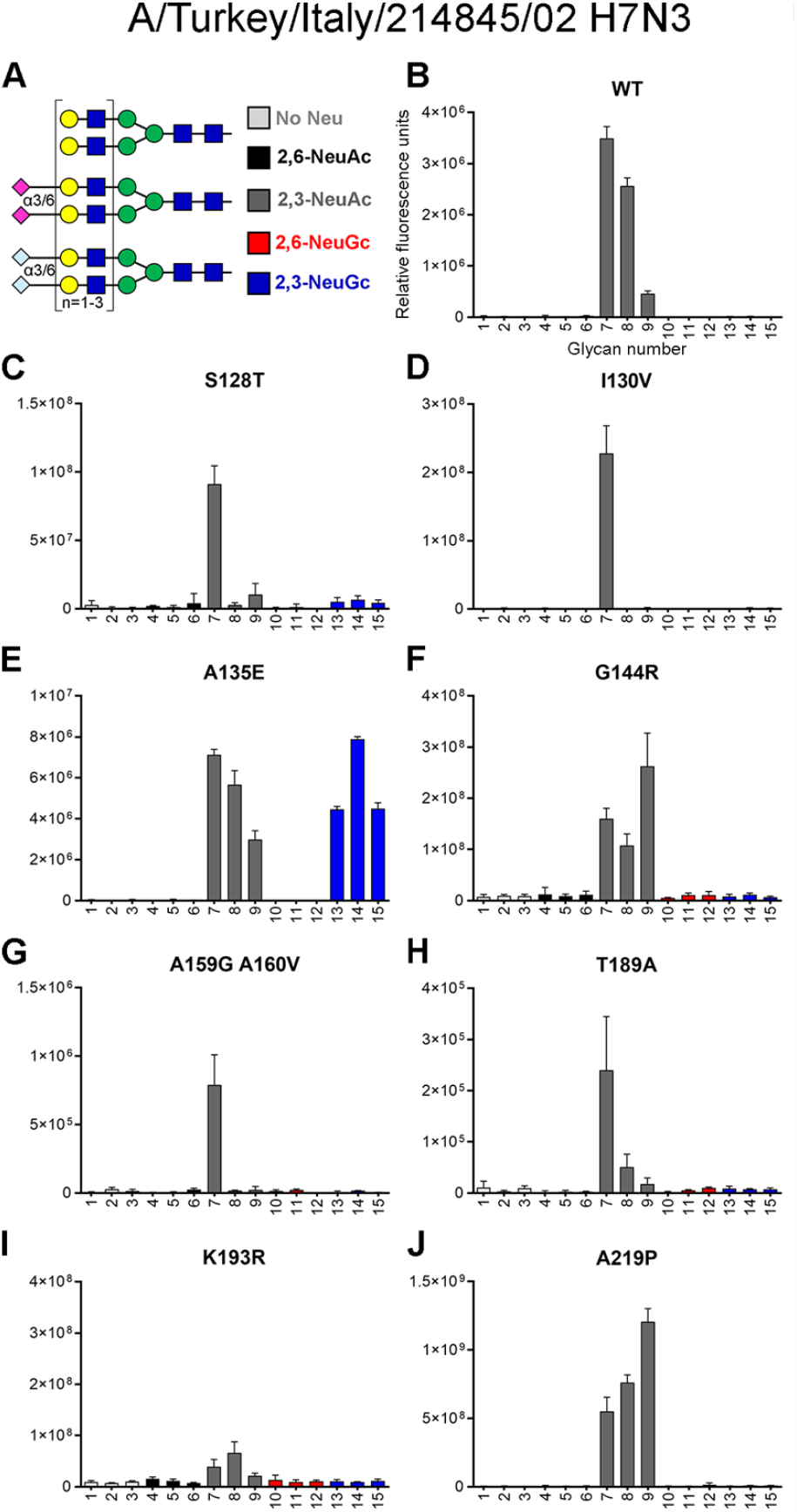
Evaluation of the binding specificities of single mutants of the HA of A/Turkey/Italy/214845/03 H7N3. (**A**) Synthetic glycans printed on the microarray (n=6), either without sialic acid (structures 1-3, light gray), with α2,6-linked NeuAc (4-6, black), α2,3-linked NeuAc (7-9, dark gray), α2,6-linked NeuGc (10-12, red) or α2,3-linked NeuGc (13-15, blue). Structures 1, 4, 7, 10, and 13 contain one LacNAc repeat, while structures 2, 5, 8, 11, and 14 have two repeats and structures 3, 6, 9, 12, and 15 contain three repeats [15]. The glycan microarray, representative for two independent assays, was used to determine the receptor specificity of (**B**) H7tu wild-type (WT), (**C**) mutant S128T, (**D**) I130V, (**E**) A135E, (**F**) G144R, (**G**) A159G+A160V, (**H**) T189A, (**I**) K193R, and (**J**) A219P.

### Amino acid 135 is essential for binding N-glycolylneuraminic acid

To identify which amino acids are critical for NeuGc binding, we mutated H7tu towards H7eq at eight different locations (128, 130, 135, 144, 159+160, 189, 193, and 219). Using the previously published glycan microarray containing glycans with terminal NeuAc or NeuGc (Fig 2A, [15]), we assessed the binding specificities of the mutants.

In the 130-loop, mutations S128T and I130V did not induce clear changes in the NeuAc/NeuGc specificity (Fig 2C-D). The amino acid at position 135 of H7 HAs is naturally diverse and has been associated with the adaptation of viruses between avian species and humans during a zoonotic outbreak of an H7N9 virus [25]. We observed that mutating position 135 (A135E) caused H7tu to acquire binding to NeuGc while maintaining binding to NeuAc (Fig 2E). Residue 143 has previously been suggested to be relevant for NeuGc recognition in H3 viruses [26]. In H7eq, R144 forms salt bridges with the 130-loop residue E135, but mutation G144R alone in H7tu did not change HA binding specificity (Fig 2F). In the 150-loop, a highly conserved tyrosine is present at position 161 in all HA subtypes except H7, H10, H12, H15, H17, and H18 [22,27,28]. Previously, it was demonstrated that a Y161A mutation changes the binding properties of an H5 HA from NeuAc to NeuGc [15,27]. However, introducing Y161A in several other HA subtypes (H1, H2, and H4) did not change binding specificities (S2 Fig). We made mutations A159G and A160V simultaneously in H7tu, but unlike the Y161A mutation in H5, we did not observe NeuGc binding with this double mutation (Fig 2G). In the 190-helix, residue 189 is next to E190, which makes hydrogen bonds to the ligand, in both H7eq and H7tu (Fig 1A and 1C). Mutation T189A in H7tu did not change receptor specificity when introduced on its own (Fig 2H). Residue 193 is important for ligand recognition (Fig 1A) [21-24]. Introducing K193R in H7tu seems to abolish all binding to the glycan array (Fig 2I), even when illuminating the glycan array with higher laser powers. Despite residue 219 being very close to the 220-loop, mutation A219P did not change binding properties of H7tu from NeuAc to NeuGc (Fig 2J). In summary, while most mutations performed on H7tu did not affect binding specificity, the introduction of mutation K193R abolished glycan-binding and A135E seems to be key for binding NeuGc while maintaining binding to NeuAc.

### Various combinations of mutations switch from NeuAc to NeuGc binding

Starting from the key mutation A135E, we continued mutagenesis by adding mutations at previously stated positions (Fig 3A). Mutating more amino acids in the 130-loop, at position 128 (S128T) or 130 (I130V), appears to abolish NeuAc binding while maintaining binding to NeuGc on the glycan microarray. The combination of mutation A135E and mutation G144R, A159G+A160V, T189A, or A219P did not change binding specificity compared to mutation A135E solely since both NeuAc and NeuGc were still bound. Whereas almost all binding was abolished when introducing mutation K193R by itself (Fig 2I), adding mutation A135E restored binding to both NeuAc and NeuGc.

**Fig 3.**
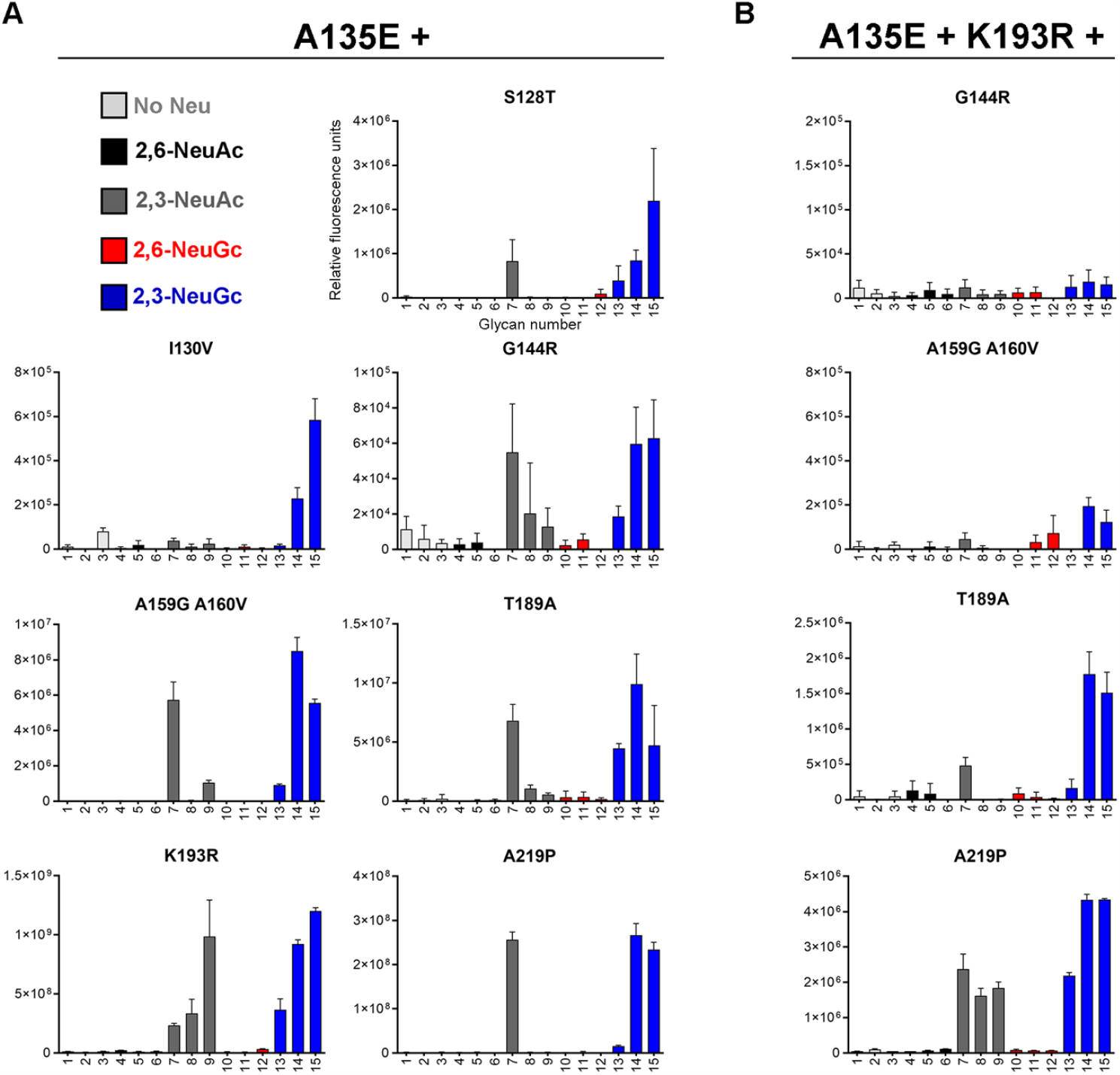
Evaluation of the binding specificities of double and triple mutants of the HA of A/Turkey/Italy/214845/03 H7N3. The glycan microarray as described in Fig 2A was used, containing glycans with terminal NeuAc or NeuGc or without sialic acid. Representative binding specificities for two independent assays for (**A**) mutant HAs containing mutation A135E and an additional mutation (S128T, I130V, G144R, A159G+A160V, T189A, K193, or A219P) and (**B**) mutant HAs containing mutations A135E, K193R, and an additional mutation (G144R, A159G+A160V, T189A, or A219P) are shown.

Since mutations A135E and K193R both affected the receptor binding properties, we further combined these two mutations with mutations that did not change binding specificity so far (Fig 3B). We found that adding mutation G144R or A159G+A160V abolished binding to the array. Adding the T189A mutation in the A135E+K193R background, switched H7tu to binding mainly NeuGc. The addition of mutation A219P did not affect the binding specificity since both NeuAc and NeuGc were still bound almost equally.

In short, we were able to modify H7tu for binding NeuGc specifically on the glycan microarray by combinations of mutations A135E+S128T, A135E+I130V, or A135E+T189A+K193R. These results show that residues in the 130-loop or 190-helix modify the specificity towards NeuGc.

### No binding specificity to avian or equine erythrocytes and tracheal epithelium observed for avian H7 mutants that bind NeuGc in the glycan microarray

A glycan microarray, as used in this study, is a sophisticated tool to investigate the binding of proteins to synthetic glycans of which we know the exact structure. However, not all glycans can be present on the array and, therefore, it is necessary to investigate the binding specificities of HAs to host cells and tissues. Therefore, we performed a hemagglutination assay with avian and equine erythrocytes and tissue staining on tracheal epithelium, which is the natural location of infection, of the same species.

While only binding NeuAc on the glycan array, the wild-type (WT) H7tu agglutinates both chicken erythrocytes, which contain only NeuAc [7,8], and horse erythrocytes, which contain mainly NeuGc [11,13,14] (Fig 4A). Therefore, a loss of binding to chicken erythrocytes would indicate a loss of NeuAc-binding. However, both types of erythrocytes were still bound by HAs with combinations of all investigated mutants (A135E, A135E+S128T, A135E+I130V, and A135E+T189A+K193R, Fig 4A) and therefore no conclusions concerning NeuGc specificity could be drawn from the hemagglutination assay. Similarly, WT and all mutants of H7tu bound both horse and chicken tracheal tissue (Fig 4C). The results demonstrate that there are some differences in NeuGc specificity between the glycan array, hemagglutination assay, and tissue staining.

**Fig 4.**
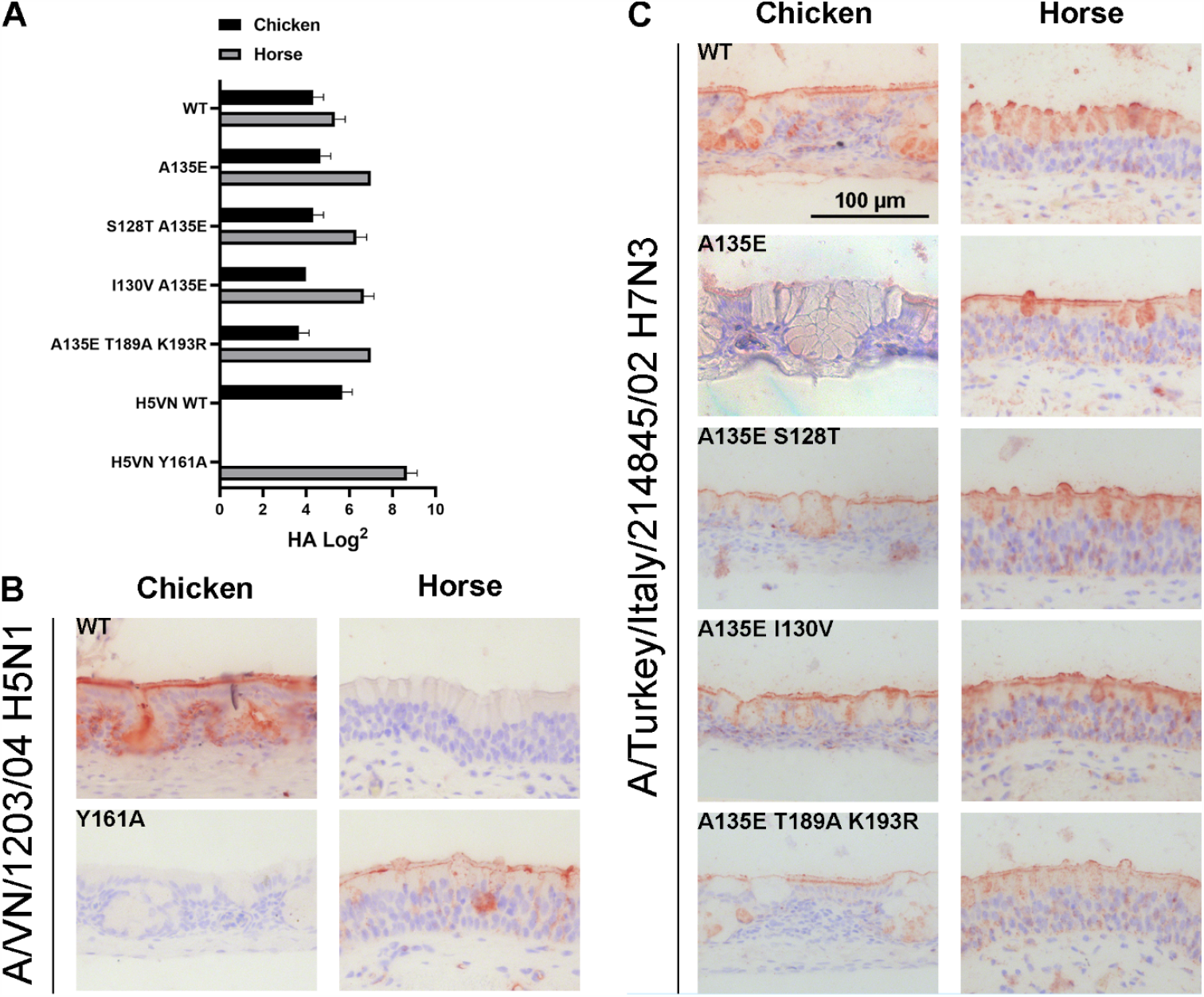
Binding specificities of (mutant) HA of A/Turkey/Italy/214845/03 H7N3 to chicken and horse erythrocytes and tracheal epithelium. (**A**) A hemagglutination assay (n=3, mean and SD shown) with chicken and horse erythrocytes was performed using H7tu WT and mutant HAs (A135E, A135E+S128T, A135E+I130V, and A135E+T189A+K193R). AEC staining is used to visualize tissue binding and images are representative for three independent assays. Tissue staining of chicken and horse tracheal epithelium is performed with (**B**) WT and Y161A mutant HA of A/Vietnam/1203/2004 H5N1 as a positive and negative control and (**C**) H7tu WT or mutant HAs as in panel A.

### NeuGc binding specificity can also be achieved in another avian H7 strain

To investigate whether the mutations that were found to switch H7tu, a virus from the Eurasian lineage, towards NeuGc binding are universal among H7 strains, we analyzed the HA of another avian strain from the North American lineage (A/Chicken/Jalisco/12283/2012 H7N3). Alignment of the HA sequences showed that the two strains differ at four amino acid positions (158, 188, 208, and 214) in the otherwise very similar RBS (Fig 5A). In glycan array analysis, similar to WT H7tu, the WT HA of A/Chicken/Jalisco/12283/12 H7N3 binds NeuAc (Fig 5B). Solely introducing mutation A135E acquires NeuGc binding and already seems to abolish some binding to NeuAc. Furthermore, NeuGc binding specificity on the glycan array was achieved by combining mutation A135E with mutations I130V or T189A+K193R. A combination of A135E and S128T resulted in a loss of glycan binding on the array.

**Fig 5.**
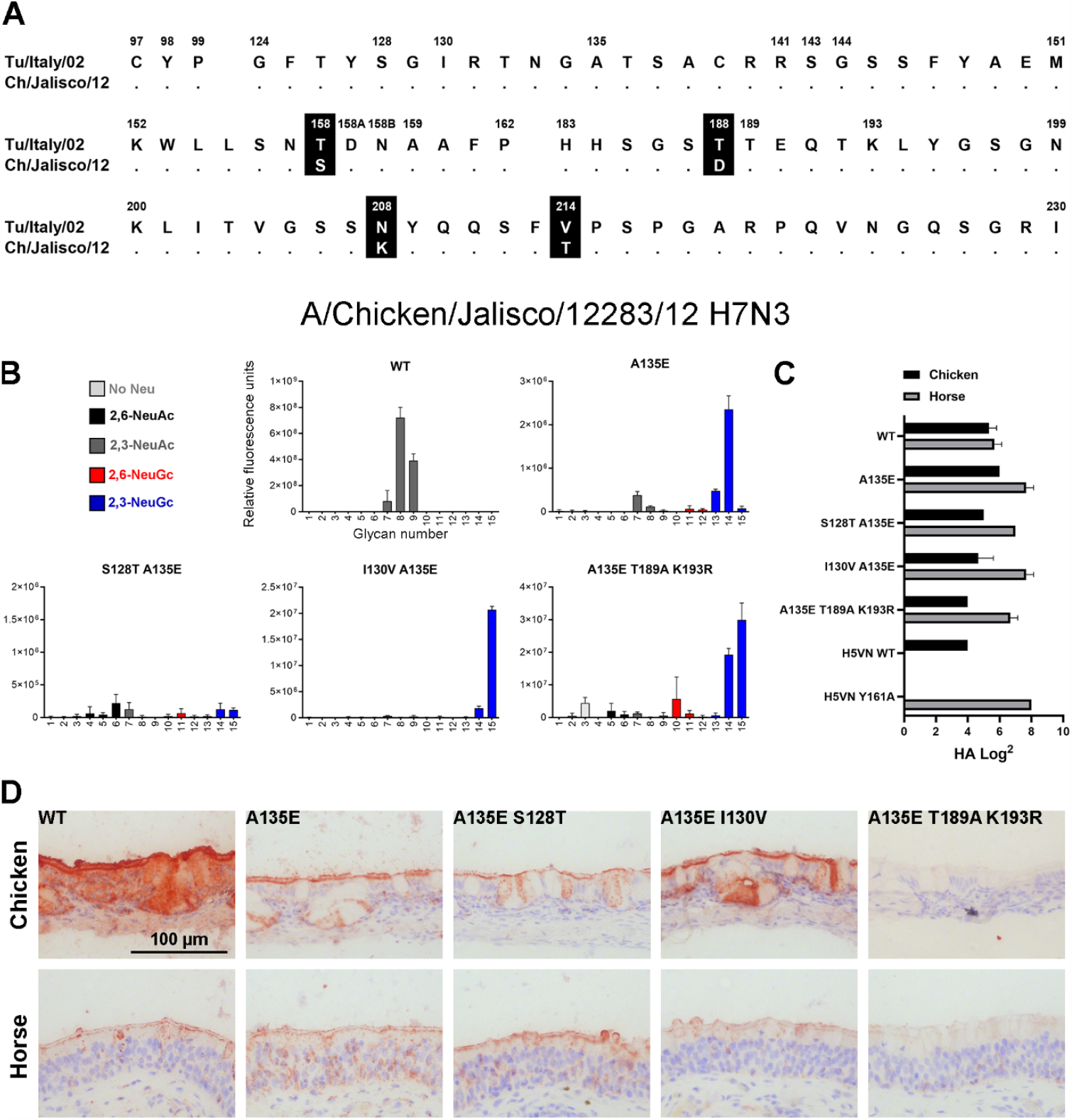
Evaluation of the binding specificities of (mutant) HA of A/Chicken/Jalisco/12283/12 H7N3. (**A**) Alignment of the RBS of the HAs of A/Turkey/Italy/214845/03 H7N3 and A/Chicken/Jalisco/12283/12 H7N3, with amino acid positions indicated above the alignment and dots indicating identical amino acids. (**B**) The binding specificities of WT HA of A/Chicken/Jalisco/12283/12 H7N3 and mutant A135E, A135E+S128T, A135E+I130V, and A135E+T189A+K193R were evaluated on the glycan microarray as described in Fig 2A. (**C**) Binding specificities of WT and mutant HAs were furthermore tested in a hemagglutination assay on chicken and horse erythrocytes (n=3, mean and SD shown). (**D**) Binding of the WT and mutant HAs to chicken and horse tracheal epithelium (controls shown in Fig 4B) is visualized using AEC staining.

The combinations of mutations A135E, A135E+S128T, and A135E+I130V did not change the binding specificity of the HA in either the hemagglutination assay using chicken or horse erythrocytes (Fig 5C) or on tissue slides containing equine and avian trachea (Fig 5D), similar to observations in H7tu. The combination of mutations A135E+T189A+K193R did not change the binding specificity in the hemagglutination assay either, but binding to both chicken and horse tracheal tissue was lost. Nevertheless, based on the glycan array analysis, we conclude that distant avian H7 HAs from different lineages can acquire NeuGc binding through identical amino acid changes.

### Equine and avian H7 strains are evolutionarily distant

The fact that avian H7 HA can be mutated towards binding NeuGc suggests that equine and avian H7 strains are phenotypically related. To investigate the genetic relationship between H7 strains, we reconstructed a maximum likelihood (ML) phylogenetic tree using HA sequences of equine H7 strains and the closest related Eurasian avian H7 strains (Fig 6A-E and S3). All equine strains cluster under a single monophyletic clade. Strains A/FPV/Dutch/1927 H7N7 and A/Fowl/Weybridge/1934 H7N7 appear to be the closest related avian strains to the equine viruses. We investigated the natural variation in amino acids at positions for which binding specificity changed (128, 130, 135, 189, and 193).

**Fig 6.**
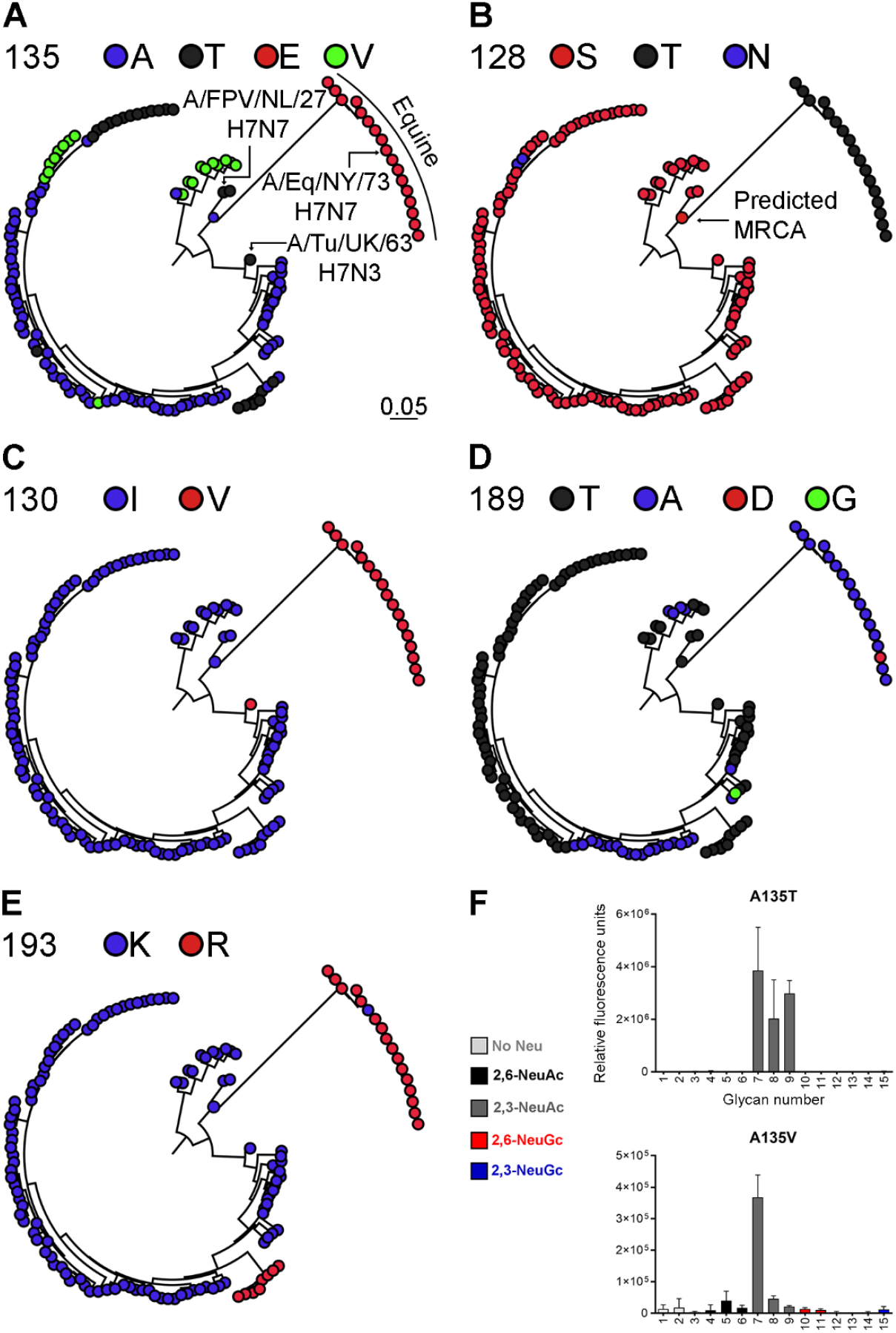
Phylogenetic tree of equine and avian H7 HA sequences and evaluation of the binding specificities of mutants of the HA of A/Turkey/Italy/214845/03 H7N3 at amino acid position 135. Phylogenetic trees of equine and avian H7 influenza A strains from the Eurasian lineage were reconstructed. The equine H7 strains cluster as a single monophyletic clade. The avian strains that are closest related to the equine strains (A/FPV/Dutch/1927 H7N7 and A/Turkey/England/1963 H7N3) are indicated, as well as A/Equine/New York/43/73 H7N7. The annotated phylogenetic tree with all strain names is displayed in S3 Fig. The variation in amino acids at positions (**A**) 135 (alanine, threonine, glutamic acid, valine), (**B**) 128 (serine, threonine, asparagine), (**C**) 130 (isoleucine, valine), (**D**) 189 (threonine, alanine, aspartic acid, glycine) and (**E**) 193 (lysine, arginine) is shown. For all positions, the amino acid of the predicted most recent common ancestor (MRCA) is shown. (**F**) Representative binding specificities on the glycan microarray (as in Fig 2A) for H7tu mutants A135T and A135V are shown.

For each selected amino acid position, we annotated the ML tree based on the variation in residues (Fig 6A-E). The predicted most recent common ancestor (MRCA) at all positions contains avian-like amino acids. Of the five investigated amino acid positions, the highest variability in amino acids is present at key position 135, although we observe a clear distinction between a glutamic acid in the equine strains and a variation of alanine, valine, and threonine in the avian strains (Fig 6A). At position 128, there is an obvious distinction between the threonine in equine strains and mainly serine in the avian strains (Fig 6B). Again, a clear difference is observed at position 130 between avian strains (isoleucine) and equine strains (valine). Surprisingly, a closely related avian strain, A/Turkey/England/1963 H7N3, also contains a valine at position 130, just like the equine strains (Fig 6C). At position 189, all but one of the equine strains contain an alanine, while there is a mixture of mainly alanine and threonine in the avian strains (Fig 6D). At position 193, nearly all equine strains contain an arginine, whereas most avian strains, except for a small clade of viruses from chickens in Pakistan, contain a lysine (Fig 6E). We conclude that there is a clear distinction between the amino acids in the equine and avian strains, with the highest variability in residues being present at position 135, which we investigated further using targeted mutagenesis.

Four different amino acids (glutamic acid, alanine, valine, and threonine) are naturally present at position 135 of avian and equine H7 viruses (Fig 6A). When the alanine is encoded by either GCG or GCA, changing a single base pair will change the amino acid to glutamic acid, valine, or threonine. We introduced all of these residues at position 135 of H7tu to investigate whether acquirement of NeuGc binding is specific for the glutamic acid. This was indeed the case, as the introduction of a threonine or a valine at position 135 did not promote binding to NeuGc (Fig 6F).

## Discussion

To elucidate the molecular determinants for NeuGc binding, we determined the crystal structure of the HA of A/Equine/New York/49/73 H7N7 in complex with its receptor analog 3′-GcLN (NeuGcα2-3Galβ1-4GlcNAc). The overall RBS structures of H7eq and A/Turkey/Italy/214845/02 H7N3 were shown to be similar. To examine the critical amino acids for NeuGc binding, we performed mutational analysis on two distant avian H7 HAs that bind NeuAc. Previously, we have demonstrated that HAs can bind either NeuAc or NeuGc [15]. Here, we demonstrate that HAs can bind both NeuAc and NeuGc, by the introduction of A135E in H7tu. The combination of mutations A135E+T189A+K193R, A135E+S128T, and A135E+I130V switched two avian HAs to bind NeuGc with some residual binding to α2,3-linked NeuAc, as determined on the glycan microarray.

We previously studied the NeuAc-specific HA of A/Vietnam/1203/2004 H5N1 (H5VN) and its Y161A mutant that is specific for NeuGc, which showed complete specificity on the glycan microarray, in the hemagglutination assay [15], and on tracheal epithelium tissue (Fig 4C). Similarly, we found that WT H1, H2, and H4 HAs bind chicken, but not horse, erythrocytes in the hemagglutination assay (S2 Fig). Contrarily, we observed that WT avian H7 HAs bound both chicken (NeuAc [7,8]) and horse (mainly NeuGc [11,13,14]) erythrocytes and tracheal tissue while binding specifically to NeuAc on the glycan microarray. This distinguishes these H7 viruses from other subtypes of influenza A. Possibly, the presence of NeuAc on horse erythrocytes and tracheal tissue, although estimated to be less than 10% of the total sialic acids [11,13,14], is sufficient to be bound by the WT avian H7 HAs. Additionally, the residual NeuAc-binding of the mutant avian H7 HAs may explain the binding to chicken erythrocytes and tissue. Another important point is that not all compounds that are present in the host are represented on the glycan microarray. Therefore, missing glycans on the array may explain the binding of avian H7 HAs to horse and chicken erythrocytes and tissue.

Besides the glycans currently present on the array (Fig 2A), glycans can be further elongated, either symmetrically or asymmetrically, be tri- or tetra-antennary, and contain one or multiple sialic acids. The addition of fucose at different positions on LacNAc structures give rise to different Lewis antigens and additional sulfate or O-acetyl groups can be present, adding another layer of complexity. Fucosylated (Lewis X) and sulfated glycans are present in the human lung [29,30] and sulfated glycans have been observed on porcine lungs [31]. For equine and avian species, the glycans in the respiratory tract have not been studied in detail yet. One study describes the presence of sialyl Lewis X structures in the respiratory tract of chickens [32]. Little is known about the glycans present on erythrocytes of different species, apart from two studies that describe that very few glycans with fucoses are present on chicken and mouse erythrocytes [33,34]. Influenza A viruses of different subtypes (H1, H3, H4, H5, H6, H7, H9, H13, H14) are known that (specifically) bind, or do not bind at all, to fucosylated and sulfated glycans [35-42]. Most relevant, avian, human, and seal H7 HAs also prefer to bind sulfated sialyl Lewis X structures [16,36,39]. In conclusion, fine receptor binding specificity regarding fucosylation and sulfation is observed in many different influenza A viruses and it could well be that the viruses used in this study also have additional receptor binding preferences, besides the distinction between NeuAc and NeuGc.

It has been suggested that recognition of NeuGc by Influenza A viruses is essential for viral replication in horses [11]. The most prevalent influenza viruses among horses currently and in the past are H3N8 and H7N7 viruses [19]. While equine H7 viruses have been shown to prefer binding to NeuGc [15,16], equine H3 viruses bind to NeuAc [16]. H3N8 viruses from horses often infect dogs [19,43], which are not able to make glycans containing NeuGc due to lack of a functional CMAH. Thus, it appears to be advantageous for H3 viruses to maintain NeuAc specificity while circulating in horses.

Equine and avian H7 strains are estimated, by phylogenetic analysis, to have separated in the mid to late 1800s [44]. Nevertheless, we demonstrated that avian H7 HAs, although genetically distinct from equine H7 viruses, are able to bind NeuGc after the introduction of two to three mutations, suggesting that these viruses are phenotypically related. The mutations converting to NeuGc binding were shown to be universal in avian H7 influenza A viruses from both the Eurasian and Northern American lineage. It would be interesting to investigate whether these same mutations induce NeuGc binding in other subtypes of group 2 influenza A viruses as well.

## Material and Methods

### Expression, crystallization, and structural determination of the equine H7 HA in complex with receptor analog 3′-GcLN

The HA ectodomain of A/equine/NY/49/73/H7N7 (GenBank ID LC414434) was cloned and expressed as described previously [15]. Briefly, cDNA corresponding to residues 11 to 327 of HA1 and 1 to 179 of HA2 (H3 numbering) was cloned into a pFastbac vector. The HA was expressed in Hi5 insect cells as described [45], after which it was purified, the trimerization domain and His_6_-tag were removed, and the HA was concentrated to 6 mg/ml.

Crystals of the H7eq HA were obtained at 20°C using the vapor diffusion sitting drop method against a reservoir solution containing 32% (w/v) polyethylene glycol 400 and 0.1 M CAPS at pH 10. The complex of HA protein with 3′-GcLN was obtained by soaking HA crystals in a reservoir that contained 3′-GcLN to a final concentration of 10 mM for 1 hour at 20°C. The crystals were flash cooled in liquid nitrogen, without additional cryoprotectant, before data collection at the Advanced Photon Source (APS) (S1 Table). Data integration and scaling were performed using HKL2000 [46]. The molecular replacement method using Phaser [47] was used to solve the H7eq complex structure, for which an apo H7eq HA structure (PDB: 6N5A) was used as the search model. REFMAC5 [48] was used for structure refinement and modeling was done with COOT [49]. The final refinement statistics are outlined in S1 Table.

### Expression and purification of HA for binding studies

HA encoding cDNAs of A/Turkey/Italy/214845/02 H7N3 [20], synthesized by GenScript, and A/Chicken/Jalisco/12283/12 H7N3 (a kind gift from Florian Krammer, Mt Sinai Medical School) were cloned into the pCD5 expression vector as described previously [50,51]. The pCD5 expression vector was adapted to clone the HA-encoding cDNAs in frame with DNA sequences coding for a secretion signal sequence, the Twin-Strep (WSHPQFEKGGGSGGGSWSHPQFEK); IBA, Germany), a GCN4 trimerization domain (RMKQIEDKIEEIESKQKKIENEIARIKK), and a superfolder GFP [52] or mOrange2 [53]. Mutations in HAs were generated by site-directed mutagenesis. The HAs were purified from cell culture supernatants after expression in HEK293S GnTI(-) cells as described previously [50]. In short, transfection was performed using the pCD5 expression vectors and polyethyleneimine I. The transfection mixtures were replaced at 6 h post-transfection by 293 SFM II expression medium (Gibco), supplemented with sodium bicarbonate (3.7 g/L), Primatone RL-UF (3.0 g/L), glucose (2.0 g/L), glutaMAX (Gibco), valproic acid (0.4 g/L) and DMSO (1.5%). At 5 to 6 days after transfection, tissue culture supernatants were collected and Strep-Tactin sepharose beads (IBA, Germany) were used to purify the HA proteins according to the manufacturer’s instructions.

### Glycan microarray binding of HA

The glycan microarray as earlier presented [15] was utilized. HAs were pre-complexed with mouse anti-streptag-HRP and goat anti-mouse-Alexa555 antibodies in a 4:2:1 molar ratio respectively in 50 µL PBS with 0.1% Tween-20. The mixture was incubated on ice for 15 min and afterward incubated on the surface of the array for 90 min in a humidified chamber. Then, slides were rinsed successively with PBS-T (0.1% Tween-20), PBS, and deionized water. The arrays were dried by centrifugation and immediately scanned as described previously [15]. Processing of the six replicates was performed by removing the highest and lowest replicate and subsequently calculating the mean value and standard deviation over the four remaining replicates.

### Hemagglutination assay

Hemagglutination assays were performed with pre-complexed HAs, as described for the glycan microarray, on 1.0% erythrocytes as previously described [50] with a starting concentration of 10 μg/ml of HA. Erythrocytes were provided by the Department of Equine Sciences and the Department of Farm Animal Health of the Faculty of Veterinary Medicine, Utrecht University, the Netherlands. The blood was taken from adult animals that are in the educational program of the Faculty of Veterinary Medicine.

### Protein histochemistry

Sections of formalin-fixed, paraffin-embedded chicken (*Gallus gallus domesticus*) and equine (*Equus ferus caballus*) trachea were obtained from the Department of Veterinary Pathobiology, Faculty of Veterinary Medicine, Utrecht University, the Netherlands. Protein histochemistry was performed as previously described [54,55]. In short, tissue sections of 4 µm were deparaffinized and rehydrated, after which antigens were retrieved by heating the slides in 10 mM sodium citrate (pH 6.0) for 10 min. Endogenous peroxidase was inactivated using 1% hydrogen peroxide in MeOH for 30 min. Tissues were blocked overnight at 4°C using 3% BSA (w/v) in PBS with 0.1% Tween-20 and subsequently stained using 5 µg/ml pre-complexed HAs, as previously described for the glycan microarray. After washing in PBS, binding was visualized using 3-amino-9-ethylcarbazole (AEC) (Sigma-Aldrich, Germany) and slides were counterstained using hematoxylin.

### Phylogenetic trees

All available, high-quality HA nucleotide sequences (i.e. sequence length is >90% of full-length HA gene segment and has <1% of ambiguous base) of avian H7Nx and equine H7N7 influenza viruses dated between 1905 and 2005 from the NCBI Genbank database were downloaded (N=944). The maximum-likelihood phylogenetic tree was reconstructed using IQ-TREE [56] using the optimal nucleotide substitution model (i.e. GTR+F+R3) based on the Bayesian Information Criterion as determined by ModelFinder [57]. Ancestral sequences were reconstructed using treetime [58].

## Data deposition

The atomic coordinates and structure factors of the HA of A/equine/NY/49/73/H7N7 in complex with 3′-GcLN are being deposited in the Protein Data Bank (PDB) under accession codes xxxx.

## Acknowledgments

We would like to thank the Department of Equine Sciences and the Department of Farm Animal Health of Utrecht University for supplying erythrocytes. We thank Andrea Gröne and Hélène Verheije from the Department of Veterinary Pathobiology of Utrecht University for providing paraffin-embedded tissues.

## Supplementary figures

**S1 Table.** Data collection and refinement statistics of H7eq in complex with 3′-GcLN.

GcLN
| Data Collection |  |
| --- | --- |
| X-ray source | APS 23ID-D |
| Space group | P3 |
| Unit cell (Å) | $a = b = 112.8,$<br>$c = 130.2$ |
| Resolution (Å)<sup>a</sup> | 48.87-2.05 (2.09-2.05) |
| Unique reflections | 116,095 (5,442) |
| Redundancy<sup>a</sup> | 9.4 (5.2) |
| Average $I/\sigma(I)$<sup>a</sup> | 16.1 (1.0) |
| Completeness<sup>a</sup> | 99.5 (93.2) |
| $R_{\text{sym}}^{\text{a,b}}$ | 0.10 (0.77) |
| $R_{\text{pim}}^{\text{a,b}}$ | 0.03 (0.34) |
| $CC_{1/2}^{\text{a}}$ | 0.99 (0.70) |
| No. molecules per ASU<sup>c</sup> | 3 |
| Refinement |  |
| Resolution (Å)<sup>a</sup> | 48.87-2.05 |
| Reflections in refinement | 110,267 |
| Refined residues | 1,455 |
| Refined waters | 688 |
| $R_{\text{cryst}}^{\text{d}}$ | 0.224 |
| $R_{\text{free}}^{\text{e}}$ | 0.251 |
| $B$-values (Å<sup>2</sup>) | |
| Protein | 70 |
| RBS subdomain (Residues |  |
| 117-265 of HA1) of chain A, C, E | 33, 40, 64 |
| Ligand of chain A, C, E | 30, 53, 100 |
| Waters | 48 |
| Wilson $B$-values (Å<sup>2</sup>) | 28 |
| Ramachandran values (%)<sup>f</sup> | 96.2, 0 |
| r.m.s.d. bond (Å) | 0.005 |
| r.m.s.d. angle (deg.) | 1.36 |
| PDB code | XXXX |

<sup>a</sup> Parentheses denote outer-shell statistics.
<sup>b</sup> $R_{\text{sym}} = \sum_{hkl} \sum_i |I_{hkl,i} - \langle I_{hkl} \rangle| / \sum_{hkl} \sum_i I_{hkl,i}$ and $R_{\text{pim}} = \sum_{hkl} [1/(N-1)]^{1/2} \sum_i |I_{hkl,i} - \langle I_{hkl} \rangle| / \sum_{hkl} \sum_i I_{hkl,i}$ , where $I_{hkl,i}$ is the scaled intensity of the $i^{\text{th}}$ measurement of reflection $h, k, l$ , $\langle I_{hkl} \rangle$ is the average intensity for that reflection, and $N$ is the redundancy. $R_{\text{pim}} = \sum_{hkl} (1/(n-1))^{1/2} \sum_i |I_{hkl,i} - \langle I_{hkl} \rangle| / \sum_{hkl} \sum_i I_{hkl,i}$ , where $n$ is the redundancy
<sup>c</sup> No. molecules for complexes refers to number of HA protomers per asymmetric unit (ASU).
<sup>d</sup> $R_{\text{cryst}} = \sum_{hkl} |F_o - F_c| / \sum_{hkl} |F_o|$ , where $F_o$ and $F_c$ are the observed and calculated structure factors.
<sup>e</sup> $R_{\text{free}}$ was calculated as for $R_{\text{cryst}}$ , but on 5% of data excluded before refinement.
<sup>f</sup> The values are the percentage of residues in the favored and outliers regions analyzed by MolProbity [59].

**S1 Fig.**
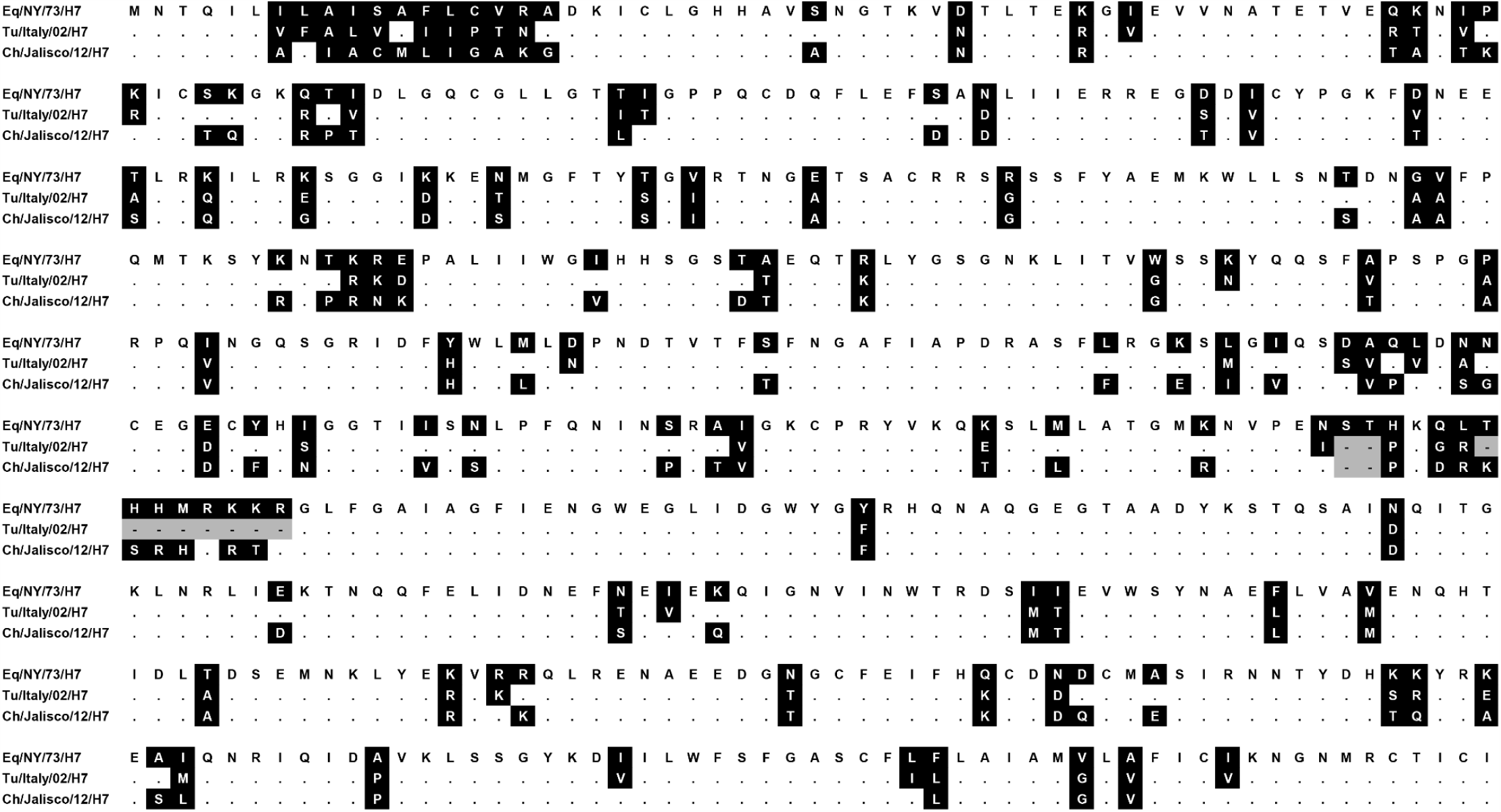
Amino acid alignment of the HAs of A/Equine/New York/49/73 H7N7, A/Turkey/Italy/214845/02 H7N3, and A/Chicken/Jalisco/12283/12 H7N3. Amino acid positions are indicated, dots indicate identical amino acids, and gray squares indicate deletions.

**S2 Fig.**
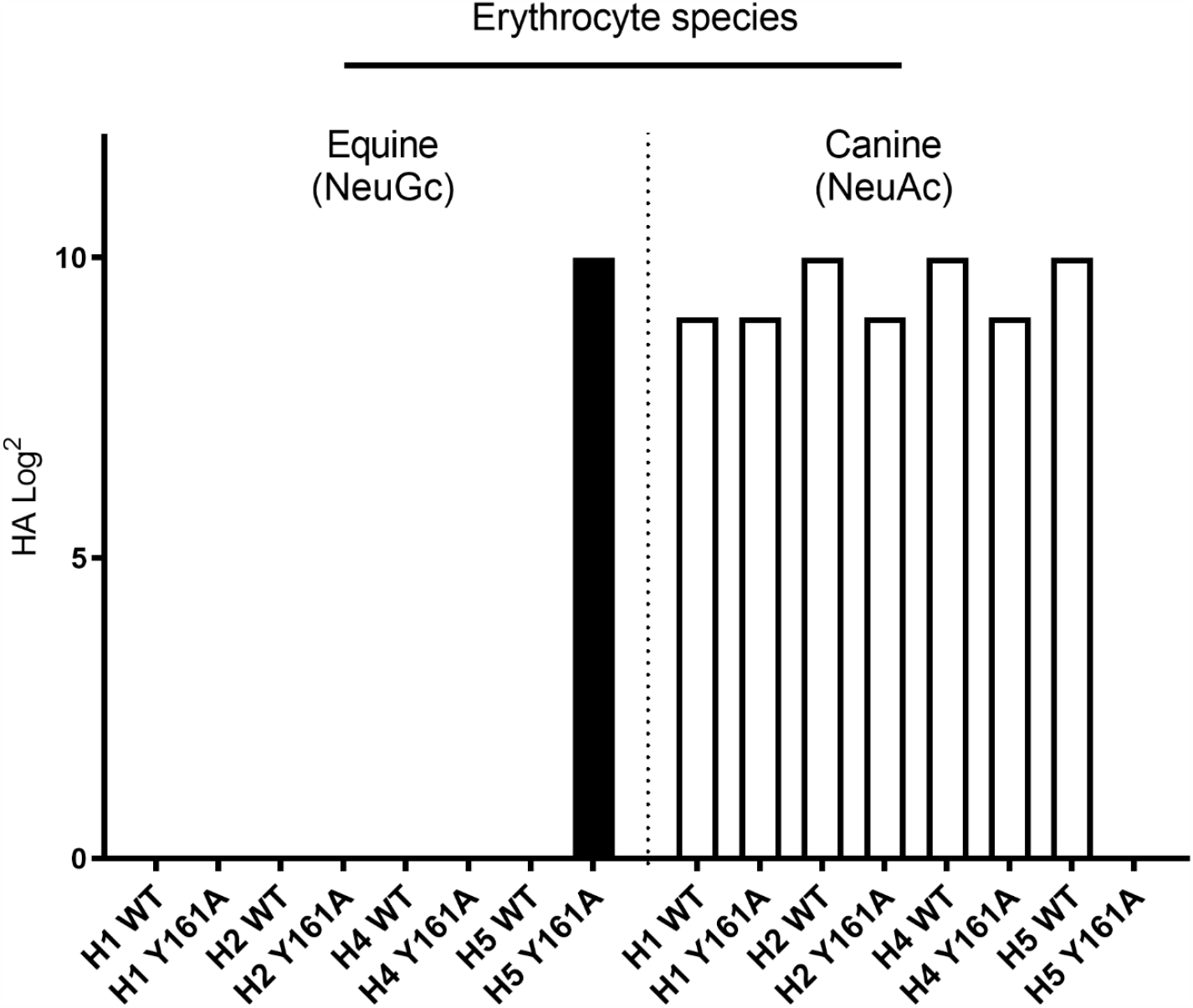
Hemagglutination assay with equine and canine erythrocytes. Equine erythrocytes contain 90% NeuGc and canine erythrocytes do not contain NeuGc. HAs of wild-type and Y161A mutants of A/Duck/Hokkaido/111/2009 H1N5, A/Duck/Hokkaido/95/2001 H2N2, A/Duck/Hokkaido/138/2007 H4N6, and A/Vietnam/1203/2004 H5N1 were investigated.

**S3 Fig.**
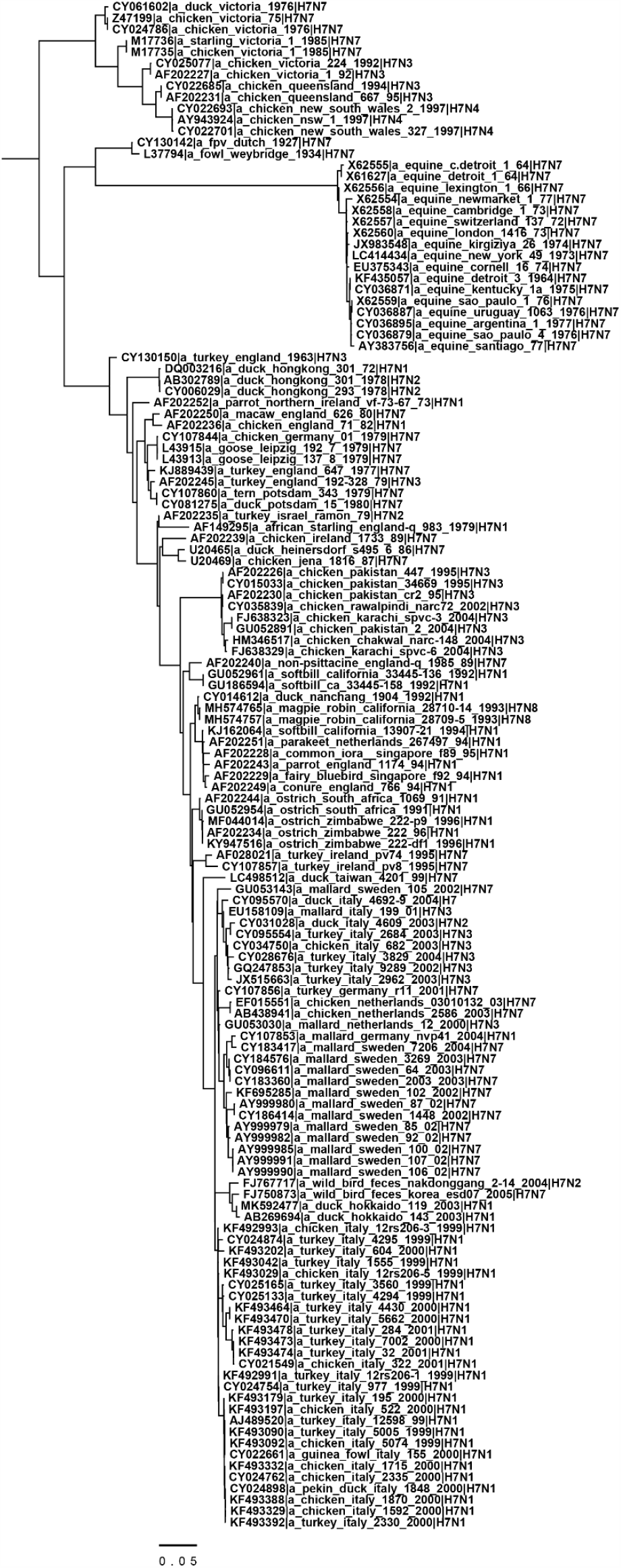
Complete annotated phylogenetic tree of equine and Eurasian avian H7 influenza A strains. The compact trees that show the variation in amino acids at positions 135, 128, 130, 189, and 193 without strain names are shown in Fig 6A-E.

